# A mathematical model of non-photochemical quenching to study short-term light memory in plants

**DOI:** 10.1101/044628

**Authors:** Anna Matuszyńska, Somayyeh Heidari, Peter Jahns, Oliver Ebenhöh

## Abstract

Plants are permanently exposed to rapidly changing environments, therefore it is evident that they had to evolve mechanisms enabling them to dynamically adapt to such fluctuations. Here we study how plants can be trained to enhance their photoprotection and elaborate on the concept of the short-term illumination memory in *Arabidopsis thaliana*. By monitoring fluorescence emission dynamics we systematically observe the extent of non-photochemical quenching (NPQ) after previous light exposure to recognise and quantify the memory effect. We propose a simplified mathematical model of photosynthesis that includes the key components required for NPQ activation, which allows us to quantify the contribution to photoprotection by those components. Due to its reduced complexity, our model can be easily applied to study similar behavioural changes in other species, which we demonstrate by adapting it to the shadow-tolerant plant *Epipremnum aureum*. Our results indicate that a basic mechanism of short-term light memory is preserved. The slow component, accumulation of zeaxanthin, accounts for the amount of memory remaining after relaxation in darkness, while the fast one, antenna protonation, increases quenching efficiency. With our combined theoretical and experimental approach we provide a unifying framework describing common principles of key photoprotective mechanisms across species in general, mathematical terms.

## 1 Introduction

Plants require light for photosynthesis, but excessive light is dangerous, because it can inflict irreparable damage to the photosynthetic apparatus. As sessile organisms, plants therefore require adaptive mechanisms to dynamically react to changing light conditions. A common strategy that has evolved in eukaryotic phototrophs [1] is the dissipation of excess absorbed energy in the form of heat, through processes collectively termed as non-photochemical quenching (NPQ) [2], which can be experimentally assessed by monitoring chlorophyll (Chl) fluorescence dynamics [3]. Analyses of the dynamics of Chl fluorescence quenching identified different NPQ components, which have been assigned to different NPQ mechanisms [4, 5, 6]: (1) the pH-regulated mechanism, qE [7], (2) the state transition mechanism, qT [8], (3) the zeaxanthin dependent quenching, qZ [4, 9], and (4) the photoinhibitory quenching, qI [10].

The discovery of the role of the xanthophyll cycle [11, 12, 13] in NPQ [14] and the identification of xanthophyll cycle mutants [15] provided a significant breakthrough in the understanding of NPQ. Since then, numerous studies supported a critical role of the xanthophyll zeaxanthin (Zx) in energy dissipation. Related to the qE mechanism, a synergistic action of Zx and the thylakoid lumen pH has been proposed [16], explaining why highest qE levels are inducible only in presence of Zx [4, 17, 18].

Based on titration studies under *in vitro* conditions, Horton and co-workers suggested that Zx shifts the pH-dependence of qE by about 1.5 pH units towards higher lumen pH [18, 19]. Furthermore, Zx was shown to modulate the kinetics of NPQ induction (faster in presence of Zx) and relaxation (slower in presence of Zx) [4, 20, 21], and was proposed to accelerate the reorganisation / aggregation of the PSII antenna [22] that accompanies qE [16, 23, 24, 25, 26]. These characteristics led to the development of the 4 state model of qE [16, 25, 27], which consistently explains the modulation of qE by the lumen pH and Zx, irrespective of the underlying quenching mechanisms and quenching sites involved in qE [24, 28].

Moreover, correlations of the Zx reconversion kinetics with the relaxation of the slower NPQ components qZ and qI indicate a function of Zx also in these processes [4, 29, 30, 31, 32, 33, 34], and Zx reconversion can be considerably down-regulated or inhibited under stress [35, 36] and photoinhibiting [36, 37] conditions. This gradual down-regulation of Zx epoxidation in response to different light stress conditions thus allows the light stress-specific adjustment of the time of Zx accumulation, and hence the modulation of the NPQ response in dependence of previous light stress, over the full time range from minutes to days or weeks [37]. It should be noted, however, that the accumulation of Zx in parallel with the activation of the different NPQ mechanisms (qE, qZ and qI) does not necessarily imply a direct role of Zx in quenching, but could simply reflect the modulation of the efficiency of quenching or a general photoprotective effect of Zx in the membrane [38]. Plants can thus store information about the illumination history to optimise their photosynthetic performance, simultaneously avoiding damaging effects of over-excitation, and Zx seems to play a crucial role for such a memory effect [28, 39, 40].

Motivated by this apparent connection between the xanthophyll cycle and NPQ induction [41], we used pulse amplitude modulated (PAM) chlorophyll fluorescence analysis to systematically investigate whether a memory of light exposure can be detected on the time-scale of minutes to hours (in contrast to other studies that investigated the illumination memory on a longer timescale [42]). We next constructed a mathematical model on the basis of our current knowledge, to provide a general description of NPQ dynamics and the associated short-term memory.

We previously argued that a major challenge of theoretical biology is to provide simple, yet robust mathematical models that are not specially tailored for one model organism but describe a variety of species [43], because only such generic cross-species models will allow discovery of common generalised principles while understanding the species-specific features. Mathematical models range in complexity from very simplified and abstract models to extremely detailed ones aiming at including all known interactions. Here, our decision on the level of the model complexity depended strongly on the specific research question that the model is designed to address.

Our aim is to find a compromise between an accurate reproduction of experimental observation and a highly reduced complexity. For this, we simplified a number of processes, which are not directly involved in the NPQ mechanism. One particular objective to derive this model is to provide a general framework, that is not specific to one organism, but can be easily adapted to different species and is specifically designed to be convenient to implement and easy to use. By providing the full documented source code, we envisage that it will be further developed by the scientific community.

The model was initially calibrated for the model organism *Arabidopsis thaliana*, a sun-tolerant higher plant. Its flexibility is demonstrated by adapting it to the non-model organism *Epipremnum aureum*, a shadow-tolerant, ornamental plant, for which measured kinetic parameters are sparse. Our model is able to realistically reproduce experimentally obtained fluorescence traces and simulate all main features of chlorophyll induction, including transient activation of NPQ [44], the dynamics of fluorescence relaxation in darkness and qualitative differences in the quenching response to different light intensities. Thus, the model serves as a tool in which the role of the main quenching components can be computationally assessed and quantified, allowing simultaneously to test existing hypotheses on short-term light memory.

## 2 Experimental Approach

Using PAM chlorophyll fluorescence analysis we quantitatively investigated the effect of short-term light memory on NPQ by comparing the fluorescence patterns of the first and second light periods. In the present study (see Fig. 1A), we examined two factors affecting the light memory in plants: intensity of incident light (varying from 100 to 900 *μ*Em^−2^s^−1^) and the relaxation time between first and second light exposure (15, 30 or 60 min).

**Figure 1:**
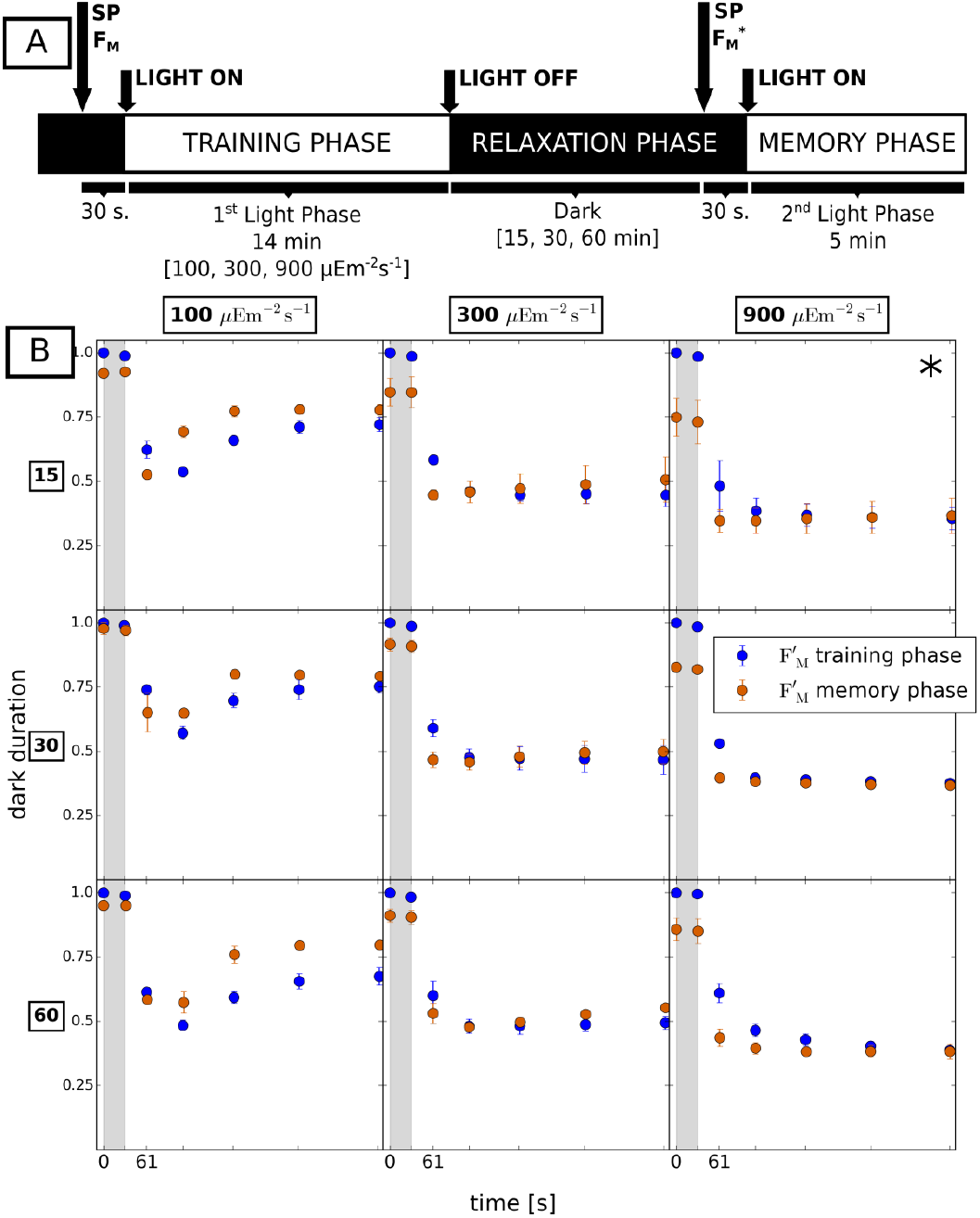
(A) Design of the experiment. Dark adapted plants were exposed to a first saturating pulse of light (SP) from which physiological parameters such as maximal fluorescence (F_M_) and photosynthetic yield (ΦPSII) were derived. 30 s into darkness a second SP was applied, then the light was switched on for a fixed (training) period of 14 minutes. SPs were applied in a defined sequence (see Fig. S1), yielding maximal fluorescence 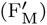. The training period was followed by a dark period, interrupted by six SP to follow the fluorescence relaxation dynamics. Subsequently, in the second (memory) period, the same illumination intensity as in the training period was applied. Each experiment was repeated three times for three light intensities (100, 300 and 900 *μ*Em^−2^s^−1^) and different relaxation times (15, 30 and 60 min). (B) Comparison of the first two 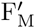 measurements taken in darkness and first five measurements in training phase (blue points) with the last two measurements taken in the relaxation phase and first five measurements in the second light phase (red points). The first measurement taken in the light (at 61 s) is lower in the memory phase than in the training phase, regardless of light intensity and time of relaxation. Error bars indicate standard deviations for three replicates except for the experiment marked with *, where the experiment was repeated eight times (see the *SI Text* for motivation).

We first analysed whether the quenching patterns differ between the two phases of light. We directly analysed the originally measured maximal fluorescence 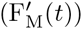 data instead of derived NPQ values (visualised in Fig. S3), to avoid mathematical distortion of the kinetics and provide more reliable information on the mechanism [45]. Fluorescence measurements are a non-invasive method for monitoring photosynthetic dynamics, providing information on the photosynthetic efficiency, protection and energy dissipation. However, each measurement can only be relative [46], and therefore at first data is normalised to the maximal fluorescence (measured after the first saturating pulse of light applied to a dark adapted plant, F_M_) and then averages and standard deviations of the three replicates are calculated. In Fig. 1B we visualise the maximal fluorescence kinetics in the first (training) and second (memory) light phase (shifted to time 0). To highlight the key features which we aimed to explain with the mathematical model, only the last two measurements taken in dark phase (marked by grey background) and the first five measurements taken in consecutive light phases are displayed (for full traces of all 22 points taken during the experiment see either the NPQ trace in Fig. S3 in the Supporting Information (*SI Text*) or extract the data from the database).

It can be observed that for all light intensities the last 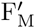 in the relaxation phase (denoted F_M_^*^) is consistently lower than in the training phase (F_M_). Likewise, the first measurement in light at 61 s shows lower fluorescence than the corresponding point in the training phase (see Tab. S1 for statistical significance). This timed response to previously experienced illumination clearly demonstrates a short-term memory. The extent of the incomplete relaxation is influenced both by light intensity and the time spent in darkness (see Fig. S4).

Based on our current understanding, these observations can be attributed to the dynamical changes in the pigment composition, especially the slow epoxidation of zeaxanthin to violaxanthin in darkness. To quantify the zeaxanthin contribution to the memory effect we measured the pigment composition at the end of each phase of the experiment (full analysis summarised on Fig. S5). Fig. 2 shows that after exposing the samples for 15 minutes to high light intensities, Zx levels significantly increased up to 50% of all xanthophyll cycle pigments (sum of violaxanthin, antheraxanthin (Ax) and zeaxanthin). Simultaneously, one hour in dark was sufficient to reduce this by half, explaining lower quenching effects in samples kept for longer periods in dark. This decrease was not as pronounced under illumination with the lowest light intensity. Moreover, zeaxanthin concentrations alone cannot explain that under 100 *μ*Em^−2^s^−1^ the fluorescence signal for the later time points is higher in the memory phase compared to the training phase. This indicates that also the relaxation of the transient NPQ depends on the light memory, a conclusion consistent with the previous findings by Johnson *et al*. [20].

**Figure 2:**
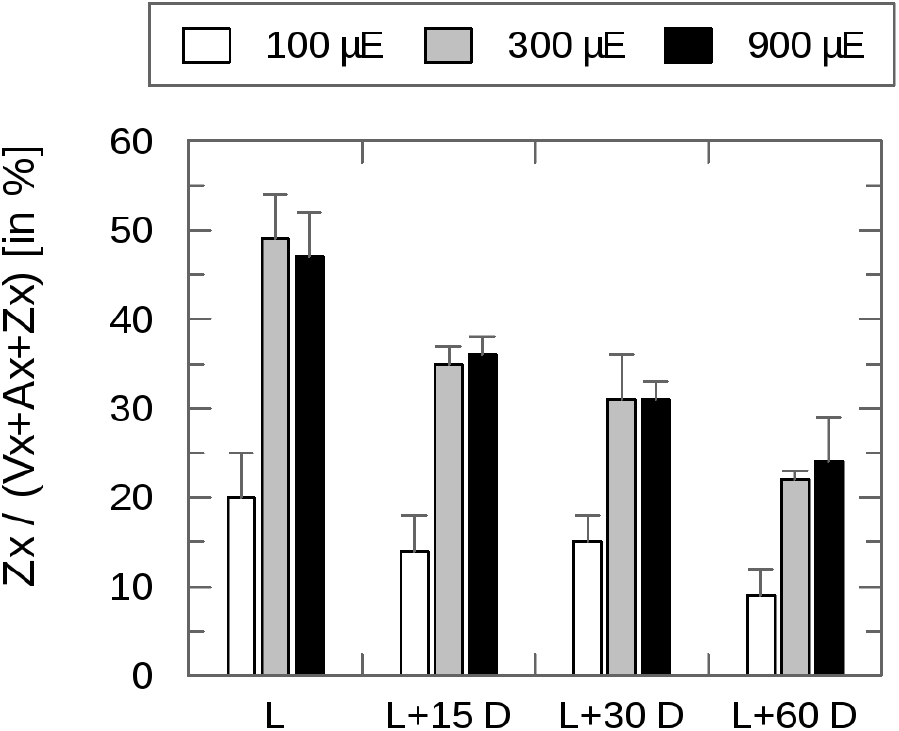
Pigment composition at the end of each phase (L=light, D=dark) presented as ratios of zeaxanthin (Zx) to total xanthophylls (Vx + Ax + Zx).

## 3 Mathematical Model

Based on our experimental results and our current understanding of NPQ, we developed a small kinetic model to verify our hypothesis on the induction of light memory and to quantify the contribution of its molecular effectors. A general schematic of the model of the electron transport chain is shown in Fig. 3. A mathematical description of the processes, as well as the source code to solve the system numerically, can be found in the *SI Text*. Since the variable chlorophyll fluorescence originates from the antennae associated with PSII [47, 48], we limit our model to the photosynthetic reactions around photosystem II, reducing the system to only six differential equations and 41 parameters (Tab. S3). Photosystem II is modelled as an oxidoreductase and is described by four possible states relating light harvesting capability with the occupation of reaction centres (following [49]). State B_0_ corresponds to the light harvesting state: reaction centres are open, ready for absorbing light energy to drive the photosynthetic electron transport chain. State B_1_ reflects the excited energy state of chlorophyll after absorption of radiant energy. From this state, in a time scale of nanoseconds, the chlorophyll can relax back to its ground state only through one of the three ways, by fluorescence emission (F), by heat dissipation (H), or by photochemistry [2] leading to state B_2_, reflecting a ‘closed’ state, in which charges are separated. State B_3_ reflects an inhibited state, in which chlorophyll is excited through absorbed light energy, but the acceptor side of the reaction centre is still occupied. Thus, charge separation and consequently photochemistry cannot occur and the excited chlorophyll can only relax to its ground state through F or H.

**Figure 3:**
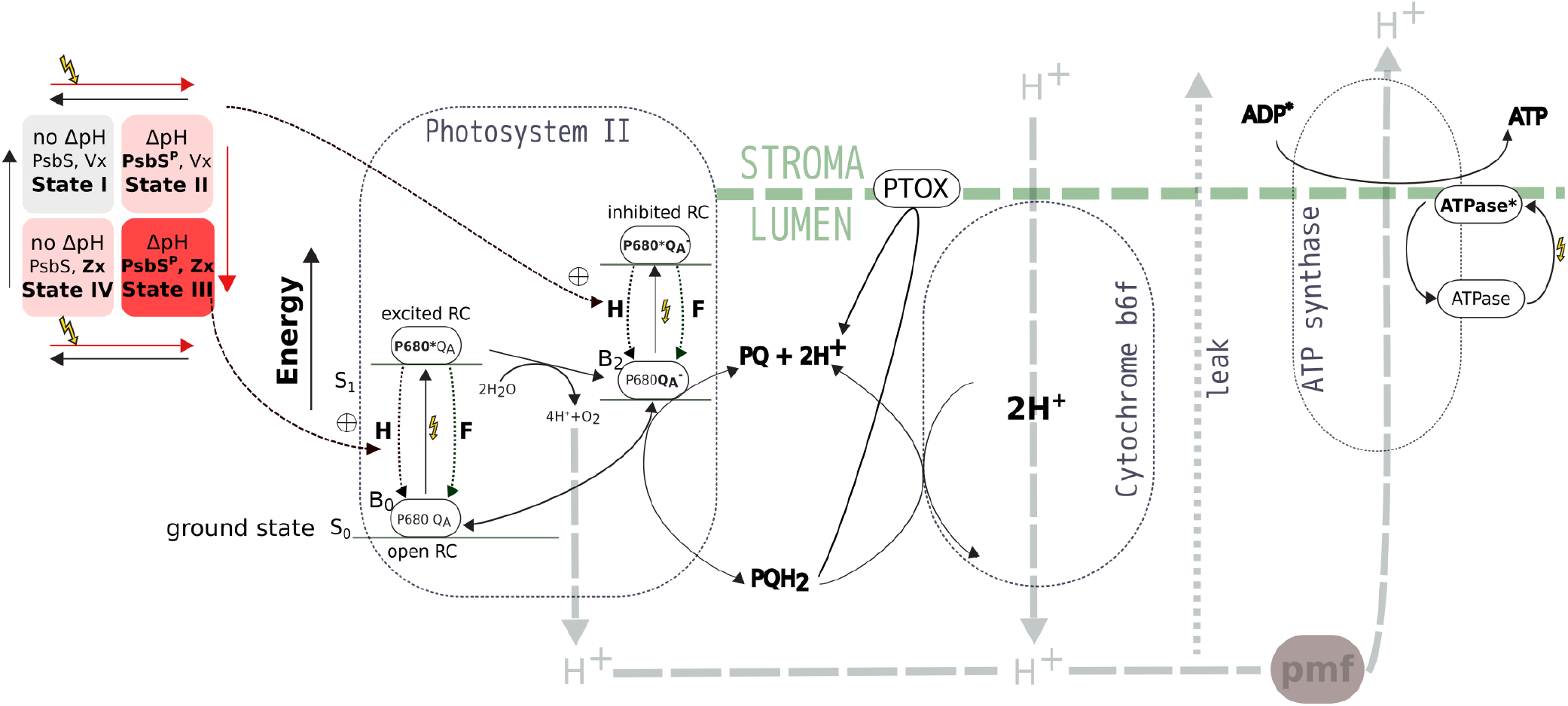
Scheme of the components included in the simplified mathematical model. It contains a detailed description of internal processes occurring inside photosystem II (for more details of the four state description of PSII see the *SI Text*) that are directly affected by the 4 state quencher. The plastoquinone pool is reduced by PSII, which also releases protons into the lumen by the oxygen evolving complex. PQ can be oxidised by cytochrome b6f and PTOX. Oxidation by cyt b6f is coupled to proton translocation from stroma to lumen. Protonation of the lumen drives the production of ATP through ATPase. ATPase is active in the light and inactive in the dark. Protons can also passively leak out of the thylakoid lumen.

Because a number of applied simplifications (see *SI Text* for justification and details) may raise concern whether the system will exhibit a biologically meaningful steady state under extreme conditions, we investigated the stationary redox state of the plastoquinone pool (PQ) over different light intensities. Under a wide range of light intensities the redox poise is maintained, meaning that the plastoquinone pool is neither too oxidised, nor too reduced to limit electron transport (see Fig. S6) [50].

### 3.1 Quencher requirements

The high-energy-state quenching represents the main component of NPQ and is induced on a time-scale of seconds to minutes [39]. Several factors are known to contribute to NPQ induction. A fast one requires the generation of a proton gradient (ΔpH) in the thylakoid membrane and a slow one is activated by low lumen pH. The quencher mechanism implemented in the model is based on a 4 state model [16, 25, 27, 51], where the fully relaxed dark state (state I) is characterised by maximal Vx concentration and de-protonated PsbS protein. In high light, a proton gradient is rapidly established, and PsbS acts as a proton sensor and thus activates quenching, constituting the fast quenching component (state II). Further reduction of the lumen pH (to 5.8) triggers de-epoxidation of Vx to Zx, leading to state III of a fully activated NPQ. From this, a transfer to darkness results in relaxation of ΔpH, and concomitant de-protonation of PsbS, leading to state IV, which still contributes to the overall quencher activity because of the slow relaxation of Zx (see Fig. 3).

These considerations lead to the overall equation for the quencher activity:

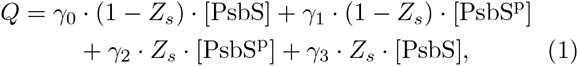

where 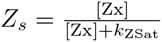 reflects the contribution of Zx to the quencher and *k*_ZSat_ is a half-saturation constant. The overall concentration of xanthophylls ([Vx] + [Zx]) is assumed to be constant and the temporal changes of [Vx] and [PsbS] are determined by appropriate differential equations (described in the *SI Text*, Eq. 9 and 10). Thus, individual and combined effects of both players on the quenching dynamics could be quantified.

The *γ* parameters were fitted to the fluorescence traces and their effect on the steady state and quencher activity was extensively studied using metabolic control analysis [52] (Fig. S7). The parameter *γ*_0_ describes the baseline quenching that does not require activation (and therefore is also active in a dark adapted sample) and was included to account for a small quenching observed in double mutants, where both PsbS and zeaxanthin dependent activation is removed [4]. The parameter was fitted to reproduce the double mutant behaviour.

### 3.2 Transiently generated NPQ

It was observed that illuminating dark-adapted plants with nonsaturating light transiently induces a strong nonphotochemical quenching [41, 47], which is explained by the transient generation of a transthylakoid pH gradient [53]. This transient NPQ relaxes within a few minutes when the ΔpH is reduced due to delayed activation of ATP and NADPH consuming reactions, including the H+-ATPase [54]. In our simplified model focusing on PSII, we therefore included a delayed activation of ATPase, by which we could realistically reproduce the overall dynamics of transient NPQ activation upon dark-light transition.

It should be noted however that also the ATP and NADPH consuming downstream reactions of the Calvin-Benson-Bassham Cycle (CBB Cycle) are redox regulated and require several minutes to achieve their full activity [55]. This leads to a similar feedback delay in ATP consumption. In a more detailed model encompassing the photosynthetic electron chain and carbon fixation, both delay mechanisms could be included and the individual contributions to the delay assessed.

For our purposes to study and explain NPQ dynamics, including one representative mechanism causing a delayed ATP consumption is sufficient. Thus, in the model ATP synthesis is mediated by an active form of ATPase (denoted ATPase*), which dynamically changes over time and directly depends on the light availability:

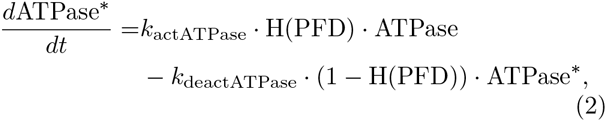

where H(x) is the Heaviside function and PFD stands for photon flux density, expressed in *μ*Em^−2^s^−1^.

### 3.3 Dynamic simulations

Our model accurately reproduces the changes in the quantum yield of chlorophyll fluorescence simulating various PAM protocols using low, moderate and high light intensities. In the model description, fluorescence is not a system variable. We therefore calculate it from the rate at which excited chlorophyll will revert to its ground state through fluorescence (*k*_F_), and not quenching (*k*_H_) or photochemistry (*k*_p_), and use the fact that the signal is proportional to the occupation of the two ground states of PSII RCs (B_0_ and B_2_ in Fig. 3) [49, 56]:

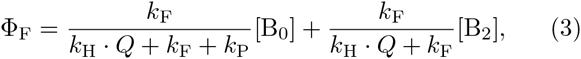

where *k*_H_ · *Q* is the rate of NPQ, modulated by the quencher activity *Q* (Eq. 1).

Determining fluorescence values from the dynamic model variables ([B_0_], [B_2_] and *Q*) using Eq. 3, allows for an *in silico* reproduction of all experimentally obtained fluorescence measurements. Fig. 4 depicts a good agreement between the simulated fluorescence traces (for continuous time) and experimentally measured 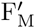 (black triangles) and F_s_ (blue triangles) values. A transient drop in fluorescence in the training phase is accurately reproduced by the simulation. Also, the fluorescence peaks in saturating light 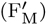 are reproduced extremely well, especially the fact that fluorescence does not fully recover after dark relaxation (*cf*. data points in the shaded regions in Fig. 1B). However, some quantitative discrepancies between the simulation and model can be observed. For instance, the first point in the light in the memory phase (Fig. 1A) is higher in the simulation than in the experiment. Nevertheless, we managed to reproduce the qualitative drop between the corresponding points, as simulated 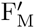 in the memory phase is still lower than the corresponding simulated 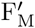 in the training phase. The steady-state fluorescence values F_s_ show discrepancies especially in the low and intermediate light intensities, but are very well captured for high light and in dark periods. One possible explanation for the deviations of model results from experimental data is that our model does not include other photoprotective mechanisms that may affect F_s_, in particular state transitions [57, 58, 59]. To study this, a more complex model is needed that is specifically designed to investigate the crosstalk and interplay between these two acclimation mechanisms. Since in the present study the focus lies on energy-dependent quenching, we consider the agreement of model and experiment as satisfactory and decide to sacrifice some degree of precision for the sake of a simple model structure.

**Figure 4:**
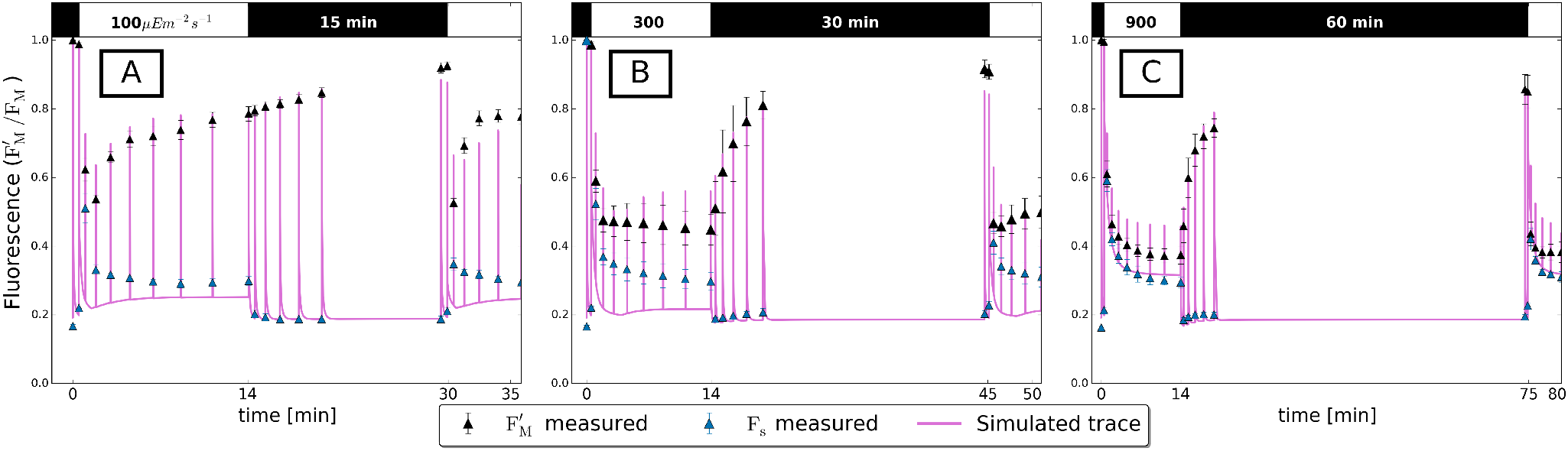
Pulse amplitude modulation traces of wild type *Arabidopsis* obtained experimentally (black triangles with error bars for 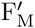, blue triangles for Fs) and simulated by the model (solid line) for three different setups. (A) Shown are the results for low light intensity 100 *μ*Em^−2^s^−1^ and shortest recovery rate of 15 minutes. The experimentally observed transient drop in fluorescence in the first minute of light exposure is qualitatively reproduced by the model. (B) Results for intermediate light intensity and intermediate relaxation phase. Under this condition neither the experiment, nor simulation show the transient drop in fluorescence. (C) The results for the highest light intensity and longest relaxation period demonstrate that the model also correctly reproduces the relaxation in darkness after one hour.

A clear benefit of computer simulations is the possibility of following the dynamics of otherwise hard to measure molecules. In Fig. 5 we demonstrate how lumenal pH changes in the response to different light intensities and how quenching components saturate under high light and relax in darkness. In Fig. 5A we plot simulated dynamics of selected internal system variables, with pH shown in the upper panel and quenching components in the bottom panel. We can observe how the transient drop in lumenal pH recovers under low and moderate light conditions but the lumen remains acidic under high light intensity. In the lower panel we can dissect the contribution to the quencher. The fast component (PsbS mediated) is quickly activated and relaxes over time, whereas the slow component continues to increase. In Fig. 5B we provide the same information in a phase phase plot, where the pH is displayed as a function of quencher activity. This representation clearly indicates the different timescales on which the system operates. Trajectories start in the dark state (low Q, pH close to 8) by rapidly reducing pH (vertical drop), before the quencher is activated (curved trajectories), and eventually the steady-state in constant illumination (red dots) is reached. The memory is visualised by the fact that the trajectories do not revert back to the initial dark state after the first dark relaxation phase, and thus the trajectories during the memory phase differ from those of the training phase.

**Figure 5:**
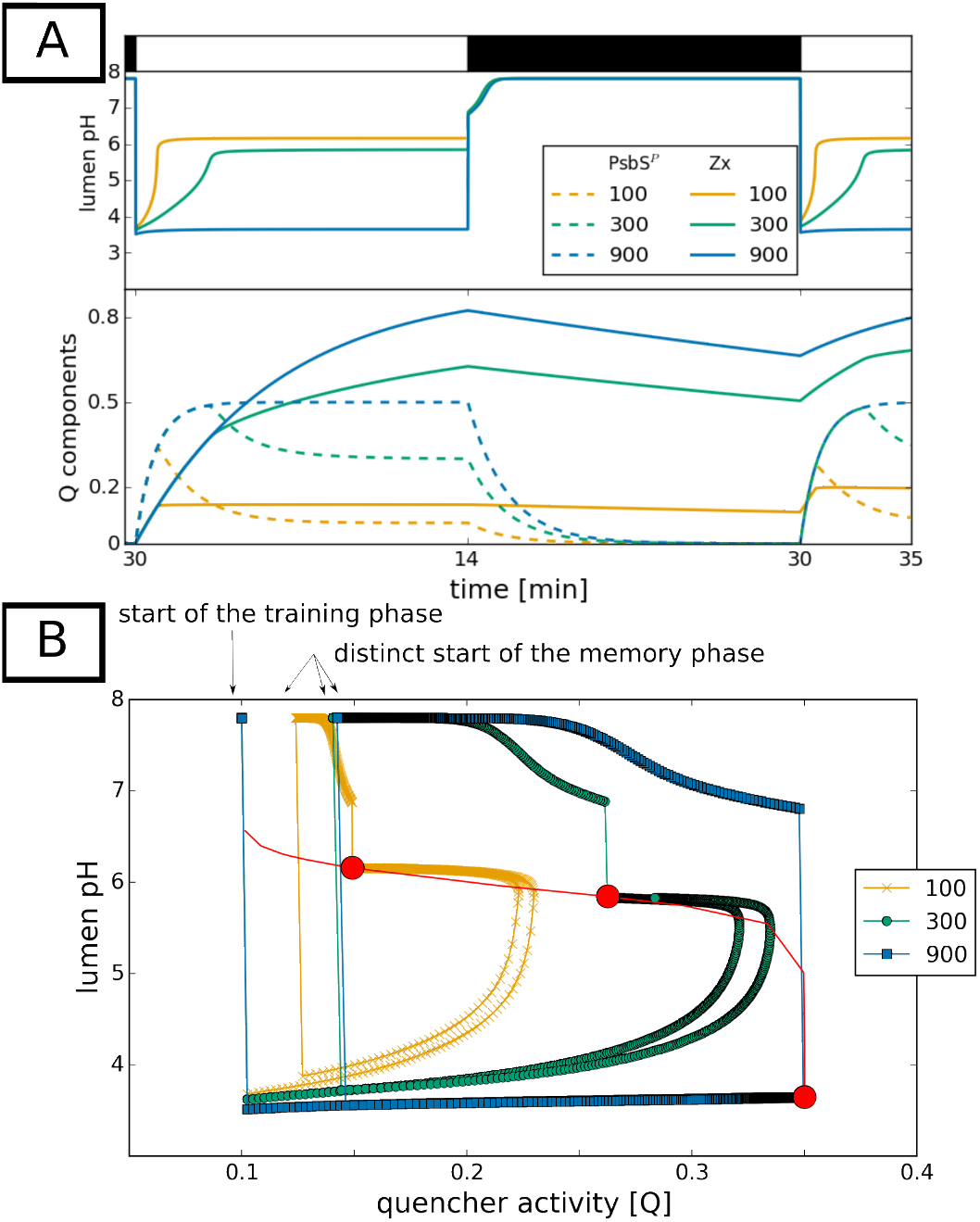
Visualisation of lumenal pH changes in the response to different light intensities. (A) Internal variables for three different light intensities. The upper panel shows how the ability of relaxing the lumenal pH is lost with increasing light intensity. The bottom panel shows the dynamics of the quenching components (solid line for the relative zeaxanthin concentration, dashed line for the ratio of protonated PsbS). (B) The phase plane trajectories of the quenching variable (Q) and the lumenal pH during the light memory protocol. The red line depicts stationary states calculated for different light intensities with three red circles marking the steady states for the three light intensities examined experimentally. The markers on the trajectories are set in regular intervals of 1 s to visualise the different time scales on which the system is operating.

With the presented mathematical model we are able to simulate different fluorescence kinetics in the training and memory phase, moreover we are able to explore which internal variables are correlated with this different dynamics and quantify their contribution.

### 3.4 Application to a non-model organism

It is a well recognised issue in the field of systems biology that modelling biological processes often requires acquiring a number of parameters [60] of which some might have a physical meaning, some may be a rough approximation of measured values, others may be fitted and some may be simply impossible to measure with current techniques. Much focus was put on developing optimization algorithms that will ease parameter estimation, but the only safe way to reduce the risk of over-fitting is to minimise the dimensions of the parameter space. One of the difficulties of using available and published kinetic models is their often huge parameter space, which makes them hard to adapt to study analogous mechanisms in other species. To demonstrate that our model is of sufficient simplicity to allow for an easy adaptation to a new and not extensively studied organism, we have adapted our model to the ornamental, shadow-tolerant plant *Epipremnum aureum*, also referred to as *Pothos*. This choice was motivated by the finding that shade-tolerant plants are characterised by longer lasting memory for leaf illumination, as compared to plants found in semi-arid climates [61].

We therefore collected the fluorescence data for *Pothos* using the same experimental setup as described above for *Arabidopsis* (see Figs. S1 and S2). The analysis of the NPQ dynamics demonstrated that under identical conditions this plant exceeds the quenching capacity of *Arabidopsis* (Fig. S3) confirming similar observations on other shadow-tolerant plants [61]. In fact, under moderate light *Pothos* already behaves like *Arabidopsis* exposed to high light. With no additional information available, we assumed that the basic principle of the protecting mechanism is the same in the two plant species, but since *Pothos* exhibits higher quenching capacity, we can reflect it by increasing the parameter *γ*_2_. We measured the chlorophyll content in both species (Tab. S2) and found a 70% higher content in *Pothos* than in *Arabidopsis*. With limited information on the electron transport chain protein abundance, we kept the same values of all internal parameters as for *Arabidopsis* and explained the more sensitive quenching response to light by increasing the factor converting photon flux density to light activation rate. With only those two changes in the parameter space we reproduced the experimentally observed fluorescence and photosynthetic yield kinetics for *Pothos* (Fig. 6).

**Figure 6:**
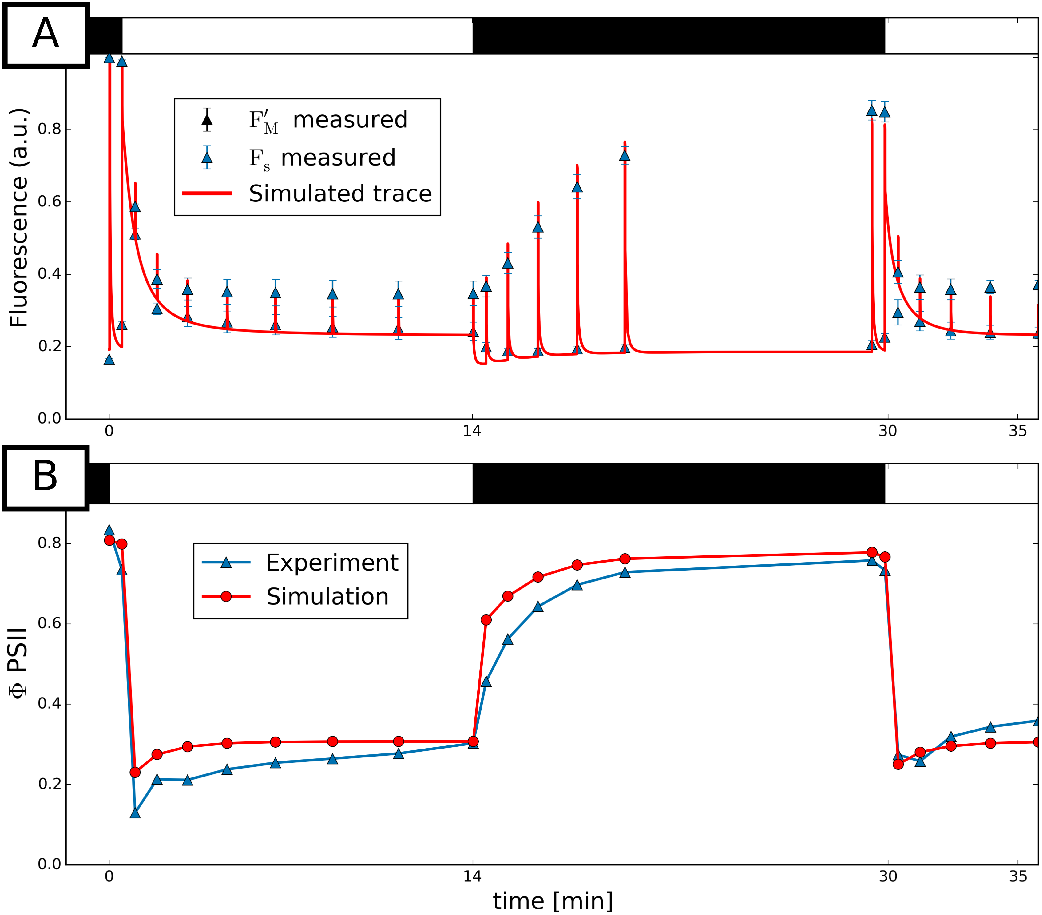
(A) Measured and simulated fluorescence traces for *Pothos* exposed to light of 100 *μ*Em^−2^s^−1^ intensity. Minimal change into the parameter space of the model enabled reproducing all key features of the fluorescence dynamics. (B) Inhibitory effect of light on the photosynthetic activity (expressed in terms of maximal yield) is reproduced in our simulation.

The agreement between the simulation and experiment allows us to suggest that the enhanced quenching capacity in *Pothos* can be explained by a more efficient energy transfer from the chlorophylls to the quencher. Possible molecular explanations might be an increased number of quenching sites or a closer spatial arrangement. To experimentally test this hypothesis detailed time-resolved fluorescence measurements would be necessary [51]. The only notable discrepancy is observed in the first measurement in light in the memory phase. However, whereas the absolute values are in slight disagreement, the variable fluorescence is accurately reproduced. This indicates, that we capture the key feature of the NPQ dynamics, but by keeping the same parameter as for *Arabidopsis*, we might slightly underestimate the protonation rate.

## 4 Results and Discussion

We have constructed a simplified mathematical model that is based on our current understanding of the quenching mechanisms. By achieving an agreement between our simulations and experimentally observed behaviour, we could confirm that this understanding is basically correct. Moreover, with the mathematical description of the process we could provide a quantitative description of the contribution of the two quenching components to light memory. It is remarkable that these findings could be obtained by a very simple model structure.

We realise that very detailed models have the potential advantage that they provide a very specific and accurate representation of experimental observations, but increasing complexity generally implies a higher level of specialisation, e. g. to one particular organism or even a single environmental condition, and moreover systematic model analyses become increasingly difficult. The virtue in model building lies exactly in the process of simplification because this process allows distilling the essential features of the investigated system which give rise to its characteristic emergent properties. Therefore, developing a simple model that can explain the same or even more than a more complex model built for the same or similar purpose, is in itself a useful scientific exercise, which generates a deeper understanding of the biological system. Moreover, simple models are easier to implement and can be generalised, but often are limited in the quantitative reproduction of the data. For instance, a model of high-energy state quenching of extreme reduced complexity aims at reproducing the key biological features of quenching [56] but is clearly limited in its capability to correctly reproduce the dynamics of quenching induction and relaxation simultaneously. On the other hand, the detailed computer model of C3 photosynthesis by Laisk *et al*. [62] was built to test whether the current understanding of photosynthesis is basically correct and is able to correctly reproduce steady state behaviour of photosynthesis and carbon fixation, but cannot reproduce NPQ kinetics. A recent model that specifically aimed at understanding the transient dynamics of NPQ, was proposed by Zaks *et al*. [63]. Employing a set of 26 nonlinear differential equations, but only considering one quenching state, the model is able to reproduce the quantitative difference in the fluorescence yield between low and high light conditions. The complexity of the latter two models makes it difficult to derive general conclusions that may be valid beyond the boundaries of single species. Further, their size makes a *de novo* implementation a very time-consuming task and the reproduction and verification of the model results is tedious, in particular since they are not provided as open source code.

We have therefore presented here a new, highly simplified mathematical model of NPQ, and employed the model not only to accurately describe the rapidly reversible components of non-photochemical quenching, such as the previously published models [56, 63], but further used it to explain the phenomenon of short-term light memory, and moreover provide a quantitative understanding of the different contributions of the well-known NPQ components.

Our model accurately simulates the changes in the fluorescence yield at low, moderate and high light intensities (Fig. 4). It further provides an explanation for the higher extent of quenching observed for plants which have previously been illuminated. Furthermore, it supports the notion that the same organisational principles of photo-protective mechanisms are present in plants as different as *Pothos* and *Arabidopsis*.

With a simple experimental setup with two light exposure periods separated by a varying relaxation time, we demonstrate that the extent of quenching does indeed depend on how long ago and how much light a plant has previously experienced – a behaviour which can safely be termed memory. We could demonstrate experimentally and theoretically that this light memory can be attributed to the slow quenching component associated with the de-epoxidation of violaxanthin to zeaxanthin triggered by low lumenal pH, which is in agreement with our current knowledge on NPQ and memory [16, 25, 27]. In the dark, epoxidation of zeaxanthin to violaxanthin is slow, so that even after 30–60 minutes the conversion is not complete. In a second exposure to light, the rapidly protonated antennae PsbS-H further contribute to quenching with an increased efficiency when zeaxanthin is still present. This conclusion is supported experimentally by direct comparison of fluorescence traces, demonstrating that light memory is affected by the length of darkness, which again determines the residual levels of zeaxanthin at the last point of darkness (Fig. 2).

Essentially, our model captures all key features of the experimental observations. However, due to the applied reduction, we cannot expect that the model will accurately describe every single aspect of NPQ dynamics, especially if processes are involved, that were not included into the model structure. For example, since the CBB Cycle is not explicitly modelled, but only summarised by a downstream consuming reaction, the effect of the redox regulation of CBB Cycle enzymes cannot be accurately reflected. Moreover, the suggested effect of Zx-induced memory on the transient NPQ in low light conditions cannot be quantitatively reproduced, because we do not incorporate any memory-dependent regulation of ATPsynthase activation. Likewise, the observed discrepancies between simulated and observed Fs levels (Fig. 4) can be explained by the fact that we do not model other acclimation mechanisms, in particular state transitions, which are known to have an effect on the steady state fluorescence [44].

Despite its simplicity, the model structure allows testing various hypotheses on the molecular mechanisms of quenching in mutants impaired in their quenching capabilities. Our simulations for the npq4 mutant, which is lacking the pigment-binding protein associated with photosystem II, but has a normal xanthophyll cycle, are in good qualitative agreement with previously published work on that mutant [64] (Fig. S8). Although mutant analysis is not in the focus of this research, this again demonstrates the flexible use and adaptability of our model and indicates its value when interpreting experimental results.

Moreover, our model can be applied to studies that focus on memory in the context of optimisation of photosynthesis (we refer readers to an excellent review covering this topic [40]). For instance, the model can be used to find a theoretical optimum size of the xantophyll cycle pool, by systematical increasing or decreasing the pool size, in order to find a balance between the benefit of a large pool leading to increased antioxidant, photoprotective activity [65], and the negative effect on the overall photosynthetic efficiency. Additionally, the response of a plant to a wide range of light protocols can be predicted, such as for example repeated light-dark cycles.

The carefully chosen selective model reduction helps to identify common underlying principles of NPQ in different photosynthetic organisms, and we expect that the model should be easily adapted even to distantly related species such as diatoms, where despite a different molecular nature of the xanthophylls, the cycle still operates according to the same principles as in higher plants [1].

Finally, thanks to the modular structure of the model, it should be a straightforward exercise to utilise this work in the context of a more detailed model of photosynthesis, such as the model on state-transitions in *Chlamydomonas reinhardtii* previously published by some of us [49], which does not include any mechanistic details of energy-dependent quenching.

## 5 Concluding remarks

We have demonstrated that our current understanding of quenching processes can be converted into more general, mathematical terms and with the implemented theory we can reproduce the most critical behavioural features of short-term illumination memory. This memory is generated by the interaction of two components of NPQ, that were previously identified by many others. The slower one, accumulation of zeaxanthin, accounts for the amount of memory lasting after relaxation in darkness, while the fast one increases the efficiency of quenching. However, our experiments do not provide evidence for an acceleration of quenching activity by previous light exposure. Rather, we propose to explain the consistently lower 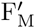 in the first seconds of the second light period by accumulation of Zx only. Therefore, plants with active short-term memory of previously experienced light initiate their photoprotection with some head-start, but at the same speed. Moreover, our computational model supports hypotheses on why shadow-tolerant plants exhibit a higher quenching capacity. Together with this manuscript we provide all necessary files to repeat and perform further experiments *in silico*, therefore we encourage our readers to treat this adaptation as an example of how our model can be used to test hypotheses regarding NPQ in other, also less studied organisms. Further, because of its simplicity and easy adaptability, the model has the potential to support knowledge transfer from lab to field. Especially in combination with cheap and easy-to-use devices to measure photosynthetic parameters outdoors (such as MultispeQ designed by the PhotosynQ project [66]), our model provides a theoretical framework in which the multitude of data can be interpreted in a sophisticated way. Thus, it can serve as a bridge to understand in how far observations obtained under controlled lab conditions allow deriving conclusions about the behaviour in real, highly fluctuating, outdoor conditions. Therefore, its real usefulness will depend on the creativity of its users.

### Plant materials and growing conditions

*Arabidopsis (Arabidopsis thaliana* ecotype Columbia 0) wild-type and *Pothos (Epipremnum aureum)* were grown in soil at the temperature of 23° C under light intensity of 90–100 *μ*Em^−2^s^−1^ with a 16 hours light/8 hours night regime. Detached leaves from three-weeks old *Arabidopsis* and two-months old *Pothos* plants were used for the measurements. To ensure perfect relaxation, plants were dark adapted for at least 12 h before the measurements.

### Fluorescence measurements

Chlorophyll fluorescence measurements were taken using PAM-2000 fluorometer (Heinz Walz, Germany) with external Halogen lamp and a DT-Cyan special interference filter (eliminating λ > 700 nm) mounted directly on the lamp. A leaf attached to the measuring window of thne fluorometer was illuminated by a train of weak probing flashes (475 nm). The average intensity of measuring light was ≤ 0.1*μ*Em^−2^s^−1^ and of saturating pulses of light was 2650 *μ*Em^−2^s^−1^. Each pulse was applied for 0.8 s and the exact times of pulse application are to be found in the *SI Text*.

### Pigment analysis

For pigment extraction, frozen plant material was homogenised in presence of cold acetone. Unsolubilised material was removed by short centrifugation and the pigment content of the acetone extract was analysed by reversed-phase HPLC according to Färber *et al*. [67].

### Simulations

The differential equations were numerically solved using the SciPy library, a Python-based ecosystem of open-source software for mathematics, science and engineering. We provide the open source code that can reproduce all figures presented in this paper, including instructions in the *SI Text*.

## Acknowledgment

This work was financially supported by the European Union Marie Curie ITN AccliPhot (PITN-GA-2012-316427) to A.M. and O.E., Deutsche Forschungsgemeinschaft Cluster of Excellence on Plant Sciences, CEPLAS (EXC 1028) to O.E. and Deutsche Forschungsgemeinschaft (JA 665/10-1) to P.J. We also wish to thank Dr Ovidiu Popa for the statistical consultations and Dr Stefano Magni and anonymous reviewers for invaluable comments that helped us to improve this manuscript.

